# Influence of ambient day light variations and age on the Iris-pupillary area ratio in beef cattle

**DOI:** 10.1101/2022.03.31.486575

**Authors:** Paulina Chojnacka, Arun HS Kumar

## Abstract

**Introduction:** Iris-pupillary area ratio (IPR) is an objective and non-invasive index of autonomic nervous system activity and stress, which can be confounded by light intensity or age of an individual. Evaluating the influence of ambient light intensity variations or age on IPR is necessary to improve the validity of IPR for its clinical application in objective assessment of welfare and stress.

**Materials and Methods:** In this study we evaluated the influence of ambient light intensity variations and age on IPR in beef cattle breeds (Limousin, Belgian Blue and Charolais) and analysed the data using correlation statistics.

**Results:** The correlation between the light intensity (20 to 500 lux; r = 0.22, p = 0.08) or age (10 to 145 months, r = 0.20, p = 0.12) and IPR was weak and statistically not significant. A sub-group analysis assessing the influence of gender on correlation between the light intensity and IPR also was not statistically significant.

**Conclusion:** Our results suggest that within the ambient light intensity (20 to 500 lux) and age (10 to 145 months) the variation in IPR is minimal. Hence our results validate the merit of IPR in objectively measuring autonomic activity/stress and demonstrate the practicality of using IPR for welfare assessment under ambient light conditions in wider age cohorts of beef cattle.

## Introduction

We have previously reported the utility of the iris-pupillary area ratio (IPR) as a non-invasive index of autonomic nervous system activity.^[1, 2]^ The sensitivity of IPR to respond to borderline changes in the activity of the para/sympathetic system allows for IPR to be used to objectively measure physiological stress levels, which is currently not feasible by any of the alternatives, such as biochemical diagnostics.^[3-6]^ The higher sensitivity of IPR is due to the ability of the iris muscles to respond to subthreshold changes in concentration of para/sympathetic system neurotransmitters.^[7-9]^ Although the specific autonomic neurons regulating the iris muscles are discrete,^[10, 11]^ physiologically the autonomic neurons function as a single network in our opinion,^[1, 2]^ hence systemic or local changes in the para/sympathetic system will be reflected by changes in the IPR.

Besides the direct regulation of IPR by the autonomic neurons, the light intensity can also influence the IPR (by regulating autonomic response), as this remains the major physiology of the iris muscles by which they help to optimise vision.^[12-15]^ Due to this confounding regulation, the clinical transition of IPR for objective measures of stress requires the demarcation of the influence of ambient light versus the degree of autonomic activity on IPR. Although changes in light intensity are known to influence IPR, the degree to which the ambient light intensity range can influence IPR is not investigated. Hence, in this study, we looked at the influence of ambient day light variation on IPR in a beef cattle herd.

## Material and methods

Digital images of beef cattle eyes were captured using a mobile phone (iPhone SE 2020). The images were taken in January 2022 at different times (morning, midday and evening) in a day over 15 days on a beef cattle farm in County Down, Northern Ireland. The beef cattle herd investigated was managed on a medium-sized farm, with around 150 animals present at the time of the study, including calves, heifers, cows, and 2 bulls. Pregnant cows and heifers, and the bulls, were kept indoors for ease of management, and cows with young calves were kept outdoors in group pens. Groups of older calves and young non-pregnant heifers were kept outdoors on pasture (grass and kale). Cattle kept indoors were fed a diet of forage (chopped silage and hay) and concentrate feed, with additional pre-calving mineral powder given to the pregnant cows and heifers. Cows kept in group pens with their calves were also fed a mixed diet of forage and concentrates, and cattle kept on pasture were provided with additional feed. Healthy and physiologically normal cattle as judged by the attending veterinary student were included in this study. As this study didn’t involve any intervention or procedures on the animals, the requirement for an ethics review was felt unnecessary.

The iris-pupillary area ratio was analysed as reported previously.^[1, 2]^ The light intensity at the time of taking the photograph was measured using the Light Meter App with the frontal camera selected. The data was analysed using GraphPad Prism software (Version 5) with significance accepted at p<0.05.

## Results

The herd consisted of three different beef breeds (Limousin, Belgian Blue and Charolais) and during the study duration the age of the subjects studied ranged from 10 to 145 months. The ambient light intensity during the duration of the study ranged from 20 to 500 lux. A total of 32 cattle (3 males and 29 females) were included in this study. The correlation (r = 0.22) between the light intensity and iris-pupillary area ratio (IPR) was weak and statistically not significant (p = 0.08), suggesting that within the ambient light intensity, the variation in IPR is minimal (Figure 1A). Similarly, the influence of age (10-145 months) on IPR was minimal, as the correlation (r = 0.20) between age and IPR was weak and statistically not significant (p = 0.12) (Figure 1B). A sub group analysis was also performed to assess the correlation of light intensity and IPR in female (r = 0.18; p = 0.19; Figure 1C) and male (r = 0.74; p = 0.09; Figure 1D) beef cattle, which showed similar outcomes.

**Figure 1:**
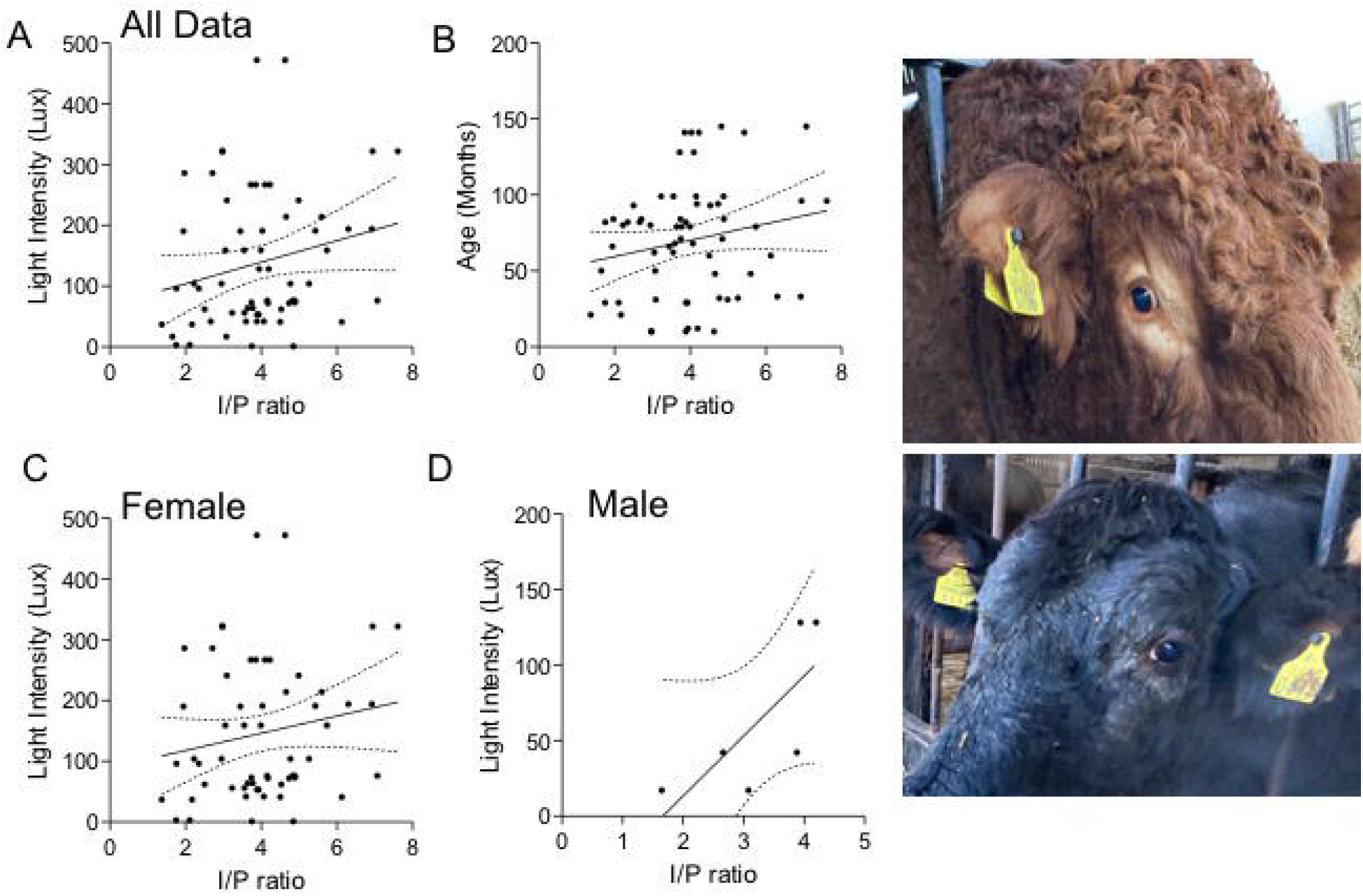
Correlation of right and left eye Iris-pupillary area ratio (IPR) with light intensity (lux) and age (months). A. Graph shows correlation (r = 0.22) of IPR with light intensity in all subjects (n=32). B. Graph shows correlation (r = 0.20) of IPR with age in all subjects (n=32). C. Graph shows correlation (r = 0.18) of IPR with light intensity in females (n=29). D. Graph shows correlation (r = 0.74) of IPR with light intensity in males (n=3). The panels on the right are two representative images used for the analysis of IPR.

## Discussion

We have previously reported the merit of the iris-pupillary area ratio (IPR) as a reliable, objective, and non-invasive index of autonomic nervous system (ANS) activity, and hence stress.^[1, 2]^ IPR is exclusively regulated by the autonomic nervous system, and can also be influenced by light intensity.^[8, 13, 14]^ However, studies evaluating the influence of the ambient light intensity under routine practical conditions on IPR are lacking. As understanding this influence of ambient light intensity on IPR will be integral to interpretation of IPR measurements, we have addressed this gap in the current study. It was indeed reassuring to observe that within a range of ambient light intensities (20 to 500 lux) in a herd of beef cattle, the influence of light intensity on IPR in healthy beef cattle was negligible. Our observations from this study further validate the merit of using IPR to reliably and objectively measure stress under routine practical conditions, as changes in IPR for any individual under ambient light intensity conditions will be clear reflection of changes in inherent ANS activity.

In this study, IPR values for beef cattle ranged from 2 to 7, which is consistent with our previous observations for unstressed individuals from different species.^[1, 2]^ This suggests the potential application of IPR measurement for objectively quantifying stress in different species, which, considering its non-invasive nature, will offer practical advantages and will prove to be beneficial for timely intervention of appropriate welfare measures. Our observations also assure that the animal husbandry practices on this particular beef cattle farm are of good standards. Based on this observation, we would also like to speculate that the baseline IPR is constant across different species and will be interesting to independently validate this in future studies.

It is well know that ANS activity changes with age.^[16-19]^ Hence, to asses if age should be factored into interpretation of IPR measurements, in this study we corelated the age with IPR. Within the age range of 10-145 months, we didn’t see any significant influence of age on IPR, which suggest to us that 1) it is not necessary to factor age into the interpretation of IPR measurements and 2) the sensitivity of pupillary muscles to autonomic neurotransmitters paraphs doesn’t decline with age. These findings further validate the potential application of IPR to reliably measure stress levels not only in a wide range of species, but also in wider age cohorts within each species. We also assessed the influence of gender on IPR in a sub-group analysis, as several reports have indicated gender differences in reflection/perception of stress, and hence autonomic activity.^[20-23]^ However, we didn’t observe any difference in the female or male cohorts with the influence of ambient light intensity on IPR. Although, our data in male beef cattle is unreliable due to a statistically insignificant sample size.

In conclusion, we have further validated the merit of IPR in objectively measuring autonomic activity/stress and have demonstrated the practicality of using IPR under ambient light conditions in wider age cohorts of beef cattle.

## Acknowledgement

Research support from University College Dublin-Seed funding/Output Based Research Support Scheme (R19862, 2019), Royal Society-UK (IES\R2\181067, 2018) and Stemcology (STGY2708, 2020) is acknowledged.

